# Gene surfing of underdominant alleles promotes formation of hybrid zones

**DOI:** 10.1101/2021.05.30.446372

**Authors:** Kimberly J. Gilbert, Antoine Moinet, Stephan Peischl

## Abstract

The distribution of genetic diversity over geographic space has long been investigated in population genetics and serves as a useful tool to understand evolution and history of populations. Within some species or across regions of contact between two species, there are instances where there is no apparent ecological determinant of sharp changes in allele frequencies or divergence. To further understand these patterns of spatial genetic structure and potential species divergence, we model the establishment of clines that occur due to the surfing of underdominant alleles during range expansions. We provide analytical approximations for the fixation probability of underdominant alleles at expansion fronts and demonstrate that gene surfing can lead to clines in 1D range expansions. We extend these results to multiple loci via a mixture of analytical theory and individual-based simulations. We study the interaction between the strength of selection against heterozygotes, migration rates, and local recombination rates on the formation of stable hybrid zones. A key result of our study is that clines created by surfing at different loci can attract each other and align after expansion, if they are sufficiently close in space and in terms of recombination distance. Our findings suggest that range expansions can set the stage for parapatric speciation due to the alignment of multiple selective clines, even in the absence of ecologically divergent selection.

## Introduction

In diploid organisms, dominance of mutations is a crucial aspect determining fitness of heterozygous individuals. Dominance can modulate the evolutionary fate of mutations (Haldane, 1924; Patwa and Wahl, 2008) and can lead to interesting phenomena, such as pseudo-overdominance (Gilbert et al., 2020) or heterosis (Yang et al., 2017). The degree of dominance is difficult to predict (Agrawal and Whitlock, 2011), and in some cases the fitness of heterozygotes is higher (overdominance) or lower (underdominance) than fitness of either homozygote. Underdominant mutations only express their deleterious effects in heterozygotes, making these mutations the simplest form of incompatibilities between different lineages. These mutations thus play an important role in maintaining stable range edges between diverged lineages via the formation of hybrid zones (Barton and Hewitt, 1985) and are also often involved in reproductive isolation and speciation (Butlin and Smadja, 2018).

It is commonly assumed that hybrid zones form as a result of (secondary) contact between two different species. However, less studied and perhaps less expected to occur, is the case of differentiation over a continuous species range, as the formation of a hybrid zone without ecological forces changing over space seems improbable. There are observed instances of such differentiation occurring, for example in systems with isolation by distance there may be observed geographic regions of distinct change in genetic makeup, such as in the greenish warbler where distinct genetic clusters within the species range have been shown to exist (Alcaide et al., 2014). In panmictic populations, Barton and De Cara (2009) have shown that underdominant mutations can combine with other pre- or postzygotic incompatibilities and may therefore play an important role in the evolution of strong reproductive isolation and speciation.

A critical question is how underdominant mutations can establish within populations despite their selective disadvantage while rare. In large populations, underdominant mutations with appreciable fitness effects should not be able to reach high frequencies because they experience a form of positive frequency-dependent selection–they are selected against while rare and positively selected only when they pass a certain frequency threshold. Wright (1941) argued that fixation of underdominant mutations is unlikely “except in a species in which there are numerous isolated populations that pass through phases of extreme reduction of numbers.” He further argued that metapopulations with local extinction and recolonization events in subpopulations are the most favorable scenario for the evolution and local fixation of underdominant mutations. This has lead to a series of influential theoretical papers investigating the role of metapopulation dynamics on the establishment of underdominant mutations, such as chromosomal inversions (Gavrilets et al., 2000; Lande, 1985). There is, however, another important and common demographic scenario that may strongly facilitate the establishment of clines in allele frequency of underdominant mutations across space: spatial range expansion.

Species range expansions are known to create a phenomenon of allele surfing, allowing for the accumulation of fixed genetic diversity at range edges (Klopfstein et al., 2006; Peischl et al., 2013). This process can impact selected and neutral variants, and has been studied extensively, particularly in regards to expansion load and deleterious variants. Another scenario less investigated, however, is the dynamics of underdominant alleles, which confer heterozygote disadvantage. During a range expansion, strong drift at the expansion front can allow underdominant mutations to cross the threshhold below which they would otherwise be selected against in their heterozygous state. A cline in allele frequencies for underdominant mutations may readily establish across the landscape. In spatially extended populations with limited gene flow, underdominant mutations can form stable clines (Barton, 1979), and may also combine with other incompatibilities (Bierne et al., 2011; Butlin and Smadja, 2018). If environmental conditions maintain multiple such clines at the same spatial location, linkage disequilibrium between the different loci will reinforce the barrier to gene flow created by underdominant mutations (Barton, 1983; Slatkin, 1975). A recent mathematical analysis has shown that even without environmental variation that keeps clines aligned spatially, clines that coincide spatially will remain together indefinitely at a stable equilibrium (Alfaro et al., 2021).

In this paper we investigate if clines in underdominant alleles can form due to allele surfing and whether clines at different loci may align in space to reinforce reproductive barriers, contributing to observed patterns of genetic diversity over space. We investigate the conditions under which range expansion can result in the establishment of such incompatibilities as well as which conditions might lead to stable hybrid zones, setting the stage for reproductive isolation and parapatric speciation.

## Methods & Results

### The model

We model the evolutionary dynamics of allele frequencies at the front of a one-dimensional range expansion, using a discrete-space serial founder event model (Slatkin and Ex-coffier, 2012). Demes are arranged as a one-dimensional array and each deme has a carrying capacity of *K* individuals. We label demes by 1, 2, 3, …, *n*. Initially, only a subset of demes is colonized, and all other demes are empty. The expansion front at time *t* is the most recently colonized deme, and we call its position *d* _*f*_ (*t*). Individuals are diploid and monoecious, and we consider a single locus with two alleles denoted *a* and *A*. Let *p* denote the frequency of the mutant allele *A* at the expansion front, that is in deme *d* _*f*_. Note that the dependence on *t* is omitted for the sake of simplicity. We consider a simple model of symmetric underdominance : the relative fitness of genotypes is given by *w*_*aa*_ = 1, *w*_*aA*_ = 1 *− s, w*_*AA*_ = 1, where 0 *< s <* 1.

A key simplifying assumption in our model is that we model the colonization of new demes as discrete founder events occurring every *T* generations (see e.g., Peischl et al., 2013, 2015). When the deme currently at the expansion front reaches carrying capacity, a propagule of size *F* is placed into the next empty deme *d* _*f*_ (*t*) + 1. The population then grows exponentially for *T* generations until the new deme’s carrying capacity is reached. The time *T* corresponds to the inverse of the expansion speed and depends on the size of the propagule and the growth rate of the population at the expansion front. The size of the propagule is determined by the dispersal rate *m* of individuals such that *F* = *Km*/2. The factor 1/2 is due to the fact that individuals migrate to each of the two neighboring demes with the same probability. During the growth phase, migration is ignored. Assuming exponential growth at rate *r* = log(*R*), where *R* is the geometric growth rate, this yields *T* = log(2/*m*)/*r* (Peischl et al., 2013). This model is a good approximation to range expansions with continuous gene flow when growth rates are larger than migration rates (Peischl et al., 2013). We also consider the case where *r* is so large that a deme grows to carrying capacity within a single generation *T* = 1, independently of the number of founders *F*.

### Fixation probability of underdominant mutations

Before we examine the serial founder event model, we consider a single panmictic population of constant size *N*. In Appendix A we use diffusion approximations to derive a formula for the fixation probability of underdominant mutations. The fixation probability of a mutation with frequency *p*_0_ at the expansion front is:

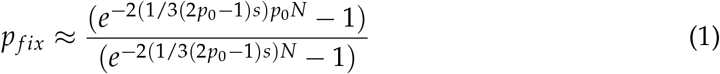

This approximation generalizes the strong selection approximation of Lande (1979) to arbitrary selection coefficients. Fig. S1 shows the fixation probability as a function of the initial frequency and for various combinations of parameters. Comparison between our analytical results and stochastic simulations shows an excellent fit between theory and simulations. Underdominant mutations are selected against as long as the initial frequency is less than 0.5, and positively selected for when initial frequencies are larger than 0.5. In the Appendix we also present an approximation for the asymmetric case where the derived allele is beneficial when homozygous, that is *w*_*AA*_ = 1 + *z > w*_*aa*_ = 1. Then the critical initial frequency at which the direction of selection changes is given by *s*/(2*s* + *z*) *<* 0.5 (see Fig. S1 B-D). This illustrates that underdominant mutations are especially prone to gene surfing because large allele frequency fluctuations at the expansion front can change the direction of selection acting on the variant and hence drive them to local fixation. A curious and related observation is that the formula for under-dominant mutations introduced as a single copy has the same shape as that of co-dominant mutations, but with an additional factor 1/3 in front of the selection coefficient. In other words, the strength of selection against rare underdominant mutations is reduced by 1/3 as compared to co-dominant deleterious mutations.

### Fixation probability at the expansion front

We next calculate the probability that a mutation goes to fixation at the expansion front in a serial founder event model. In Appendix A we argue that we can approximate the fixation probability at the expansion front by replacing *s* with *sT* and *N*_*e*_ with the number of founders *F* in equation 1. The intuitive justification for this approximation is that the action of selection over *T* generations is very similar to a single generation of selection but with a stronger selection coefficient *sT*. Furthermore, the amount of drift during founder events is the main source of stochasticity and dominates the drift during the growth phase, so we can replace *N*_*e*_ with *F*. Fig. 1 compares this approximation with individual-based simulations showing that our approximation works very well. In particular, we see that only the compound parameter *sT* matters. If *sT ≪* 1 the mutation behaves as if it is neutral and the fixation probability is approximately 1/*F*. Thus, for rapidly expanding populations (that is, small values of *T*), selection is inefficient and mutations will readily surf if they appear at the expansion front. If in addition the number of founders *F* is small, the probability for surfing becomes very large as it approaches the neutral fixation probability 1/*F*.

**Figure 1:**
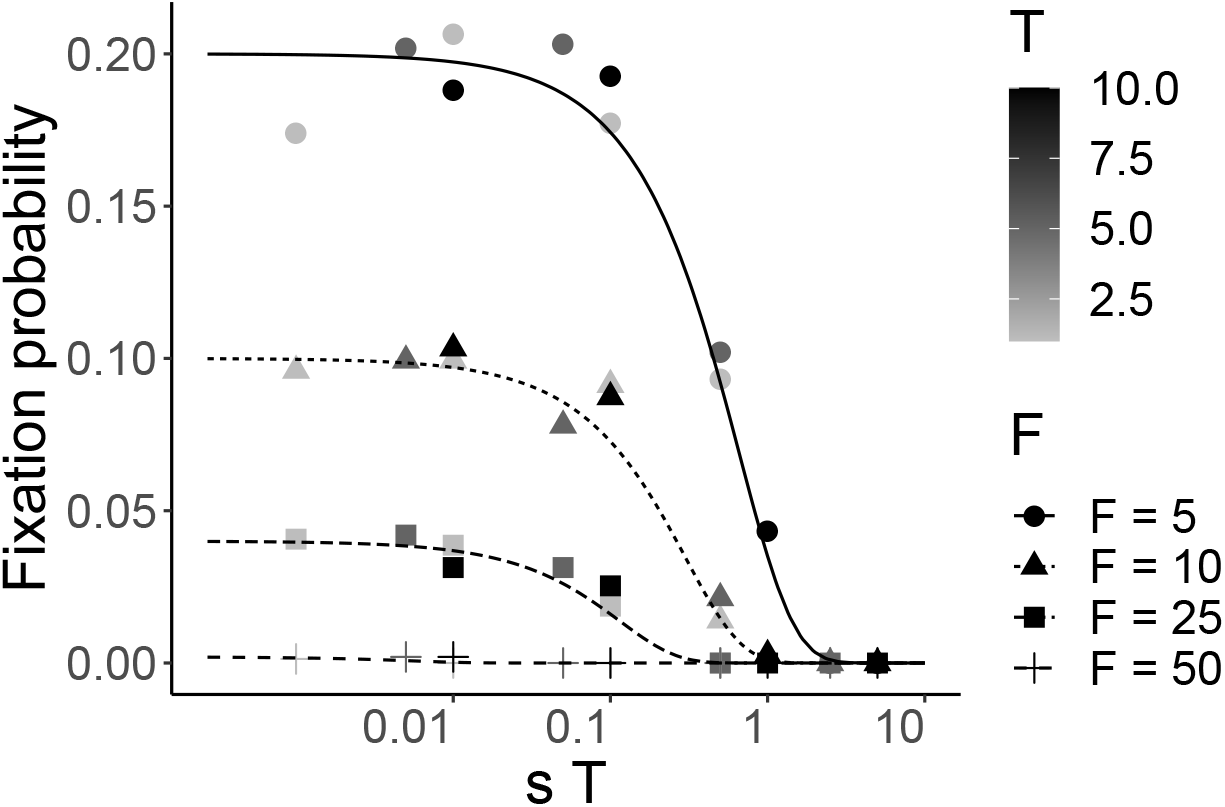
Fixation probability of an underdominant, beneficial mutation in a serial founder effect model. *F* is the number of founders, *T* the time between founding events, and *s* the strength of selection.

### Surfing can lead to stable clines in simulations

In spatially structured populations, locally fixed deleterious mutations will eventually be purged when gene flow reintroduces genotypes from locations where the mutation is not fixed. Underdominant mutations, however, can be maintained indefinitely at a stable equilibrium if they establish locally because the joint action of selection and gene flow against heterozygotes leads to stable clines (Barton and Hewitt, 1985). This has an important consequence: while the establishment of mutation load during range expansions is generally a transient phenomenon, surfing of underdominant mutations can lead to long-lasting clines and so-called hybrid zones.

We use forward-time, individual-based simulations in SLiM v.3.4 (Haller and Messer, 2019) to further investigate the formation of such clines due to gene surfing. Using a non-Wright-Fisher model (see SLiM manual), we generate a linear landscape of 250 demes with carrying capacity *K* = 25 per deme. We initialize the population in one end deme and allow expansion to proceed across empty space via a stepping stone model of migration with *m* = 0.1. These parameter combinations (high migration, small population sizes) approach a more continuous-space landscape model, and can thus be seen as complementary to our discrete-space serial founder event model. SLiM input scripts are available on github.com/kjgilbert/UnderdominantSurfing, for exact details of simulation parameters and settings. Underdominant alleles enter the population as random new mutations (genome-wide mutation rate *U* = 0.05, genome size of 10kb, and per-bp mutation rate *µ* = 5*e*^*−*7^) with heterozygotes for single locus mutants having a fitness of 0.9. The first panel of Fig. 2 shows that clines in underdominant allele frequency do establish during range expansions. Intriguingly, Fig. 2 also shows that multiple clines coincide spatially, suggesting that clines do not evolve independently but may either form simultaneously during a simultaneous surfing event, or move over space after formation until cline centers coincide for multiple loci. In the former case, if the mutant alleles surf together, it is straightforward to understand that the clines will coincide in geographic space. Simultaneous surfing requires that the mutations fall on the same genomic background or that recombination is sufficiently high so as to prevent interference between the surfing of one underdominant allele on one genomic background versus another. However, we can also see that simultaneous surfing is not always the case and that clines do move over space to eventually later coincide. From the earlier generation snapshots (lighter colors) in Fig. 3, we can see several loci which initially formed non-coinciding clines and over time moved to the same geographic location by the final generation (black lines).

**Figure 2:**
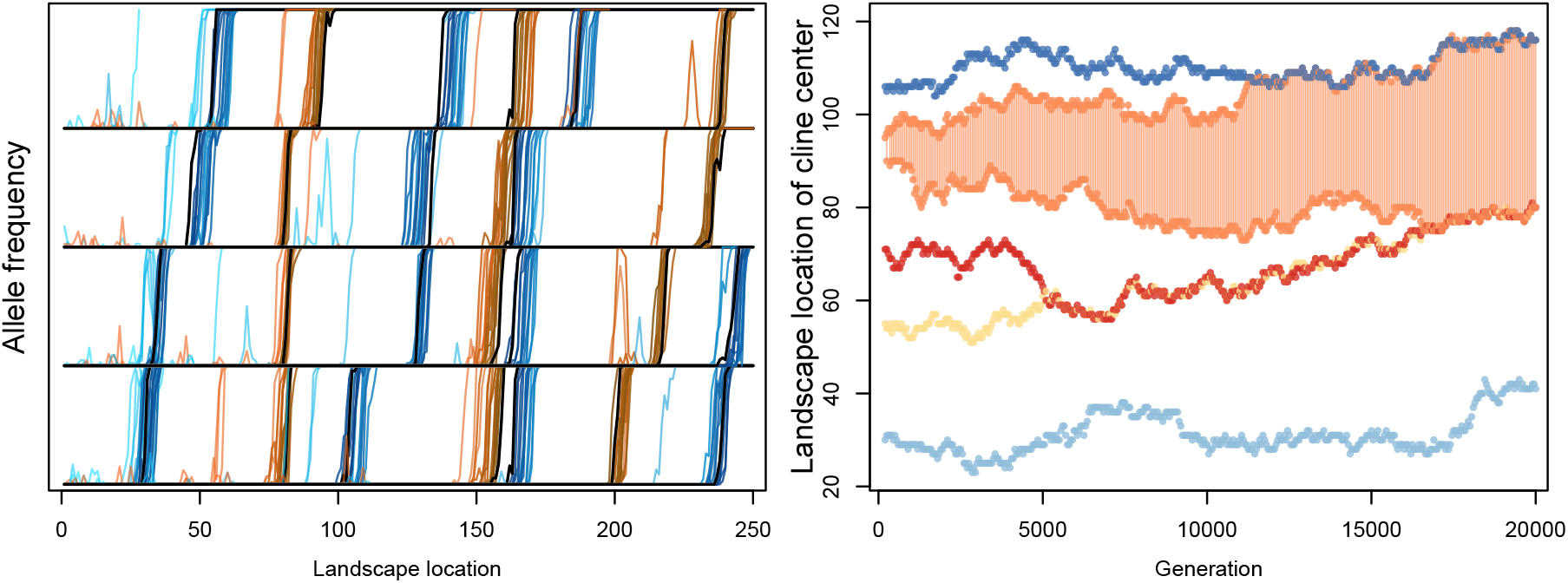
The formation (left panel) and movement (right panel) of clines for underdominant alleles during a range expansion. In the left panel, a single replicate simulation for cases with a mutation rate (see main text) is shown as an example for the formation of clines in allele frequency of underdominant mutations via allele surfing. Loci are separated onto rows only to visually distinguish different loci that have aligned in the same location on the landscape. 26 clines of underdominant mutations formed in this simulation replicate. The progression through time for each cline is shown every 100 generations from either light blue or light orange (*t* = 100) to black (*t* = 2000). In the right panel, a single replicate simulation is shown for cases with manually introduced underdominant alleles (see main text). These simulations contain only 5 loci, conditioned on each of these loci forming a cline. The movement of these clines is then examined over space for 20,000 generations to investigate the alignment of clines in space. Each color is the location of a cline center for a given locus (five colors), and the orange locus with shading between it has formed two clines, with the underdominant mutation as a fixed homozygote in the space between the two clines.

**Figure 3:**
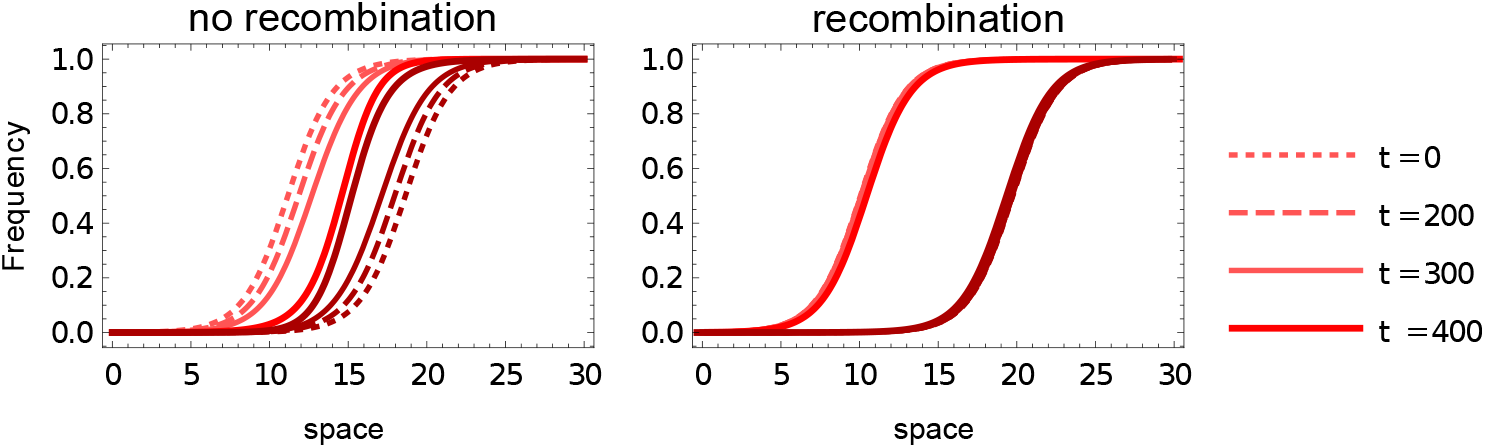
Temporal evolution of two clines at two different loci and spatial locations. Parameter values: *s* = 0.1, *d* = 0.05, and *r* = 0 left and *r* = 0.5 right.

### Clines at different loci can align spatially

To further investigate the movement and interaction of clines through space, we next conduct a set of simulations where the underdominant alleles are manually introduced into the population at a set number of generations apart, always at the range edge, for simulation efficiency. These simulations are conditioned on the mutant underdominant allele at every locus being retained in the population (*i*.*e*. simulations where an under-dominant allele is lost from the entire population are discarded and rerun). The right panel of Fig. 2 shows one replicate simulation with five such manually introduced underdominant loci that have surfed and formed six clines. The center of each of these clines is measured as the deme where allele frequency is closest to 0.5 (because clines are steep, as can be seen in the left panel of the figure, there is a +/ *−* 1 deme margin of the exact deme center), and the position of these centers is tracked over space through time. Because we want to here examine the movement of clines over space, migration was increased to *m* = 0.25 and carrying capacity per deme was instead *K* = 50. The five mutations enter the population at generations *t* = 50, 85, 120, 155 and 190, and their movement is then tracked from generation 200 onward. We observe that the clines do drift randomly over space, and when they happen to approach each other, once they reach a critical distance apart, they couple together in space and remain coupled for the remainder of the simulation (Fig. 2). We also observe that in the fourth (orange) locus, the underdominant allele has surfed only past the 0.5 allele frequency threshhold leading to its eventual local fixation behind the expansion front and a cline where it increase from frequency 0 to 1, but subsequent strong drift removes the allele at the range front creating a second cline in frequency from 1 to 0. This locus thus forms two clines in space with homozygous mutants residing between them (shaded orange area in Fig. 2). These six initial clines at five loci drift over space and eventually align at several loci resulting in three multi-locus clines across the landscape after the expansion has completed. The critical distance between cline centers before coupling of clines is quite small, given the steepness of these clines, with uncoupled cline centers never reaching closer than four demes apart, out of 20 replicate simulations of five loci each (Fig. S2).

### When do clines attract each other?

To investigate the conditions for when clines will align spatially after their formation at different loci we introduce a deterministic model (Bü rger, 2017). Our goal is to model the dynamics of clines in the wake of the expansion wave or after the expansion has ended. We therefore ignore genetic drift and use a two-locus model of migration, selection, and recombination in one-dimensional continuous space. We use the following fitness scheme for two-locus genotypes:

|  | aa | aA | AA |
| --- | --- | --- | --- |
| bb | 1 | 1-s | 1 |
| bB | 1-s | (1-s)<sup>2</sup> | 1-s |
| BB | 1 | 1-s | 1 |

We denote the frequency of haplotype *ij* by *p*_*ij*_, *i ∈ a, A, j ∈ b, B*. The marginal fitness of haplotype *ij* is given by *w*_*ij*_ = ∑_*kl*_ *w*_*ij,kl*_ *p*_*kl*_, and the mean fitness of the population is given by 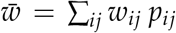. The temporal evolution of the haplotype frequencies is then given by

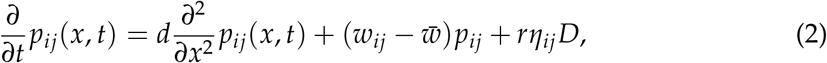

where *D* = *p*_*ab*_ *p*_*AB*_ *− p*_*aB*_ *p*_*Ab*_ is a measure for linkage disequilibrium. The parameter *d* is the diffusion term describing the variance of the displacement of offspring from the origin of their parents, and *r* measures the strength of recombination. Furthermore, *η*_*ab*_ = *η*_*AB*_ = *−*1 and *η*_*aB*_ = *η*_*Ab*_ = 1. We can simplify the system of differential equations by re-scaling its parameters. We introduce the natural spatial scale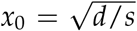 and rescale time by *τ*_0_ = 1/*s*. Dividing both sides of the equation by *s* we can write:

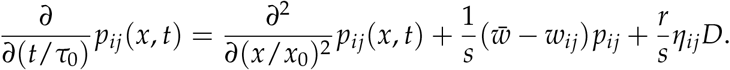

If we re-scale time and space accordingly with *t ← t*/*τ*_0_ and *x ← x*/*x*_0_ the equation becomes

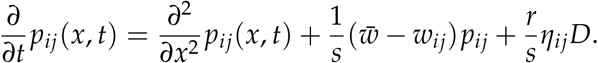

At first order, the fitness difference between the haplotype *ij* and the mean fitness is proportional to *s*, so that the second term of the right hand side is independent of *s*. For *r* = 0, the equation is thus dimensionless, and for *r >* 0 we can introduce another rescaled parameter *r*/*s* that should measure the timescale of selection and recombination.

Fig. 3 shows an example for numerical solutions of this system of partial differential equations using the original parameters. We find that the spatial width of a cline is proportional to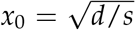, in agreement with the fact that dispersal tends to homogenize allele frequencies across the cline whereas selection tends to remove unfit heterozygotes from the population and make the cline steeper (Barton and Hewitt, 1985). We find that two clines at different loci will tend to spatially align in the absence of recombination (Fig. 3 left panel) but remain apart over the considered time period if recombination is strong (Fig. 3 right panel).

To investigate the cause for clines attracting each other and aligning after they have formed, we consider an initial state of two clines which have already formed and are a given distance apart. The first cline is a result of the surfing of allele *A* to fixation at the first locus and the second cline has formed later in the range expansion (further along the landscape) due to the surfing of allele *B* at the second locus. To understand why the clines attract each other, we can examine the distribution of allele and haplotype frequencies across the landscape. See fig. S3 for a visual depiction of these frequencies. If we examine demes over space from the origin of the expansion (left) to the conclusion of the expansion (right), we observe the following. To the left, there are populations where only the ancestral haplotype *ab* is present, and as we proceed right we pass the first cline in a *A* where we have a mixture of *ab* and *Ab*, and further right only *Ab* is present until we pass the second cline and encounter *AB*. If we focus only on the *A* cline in the absence of recombination and follow what happens through subsequent generations, we observe that initially, migrants with genotype *abab* come from the left of the cline, and *AbAb* from the right. These genotypes are equally fit, and the frequency of the allele *a* is maintained at an equilibrium value due to migration-selection balance, creating a stable cline in space. However, heterozygous migrants with genotype *ABAb* will eventually arrive from the *B* cline further to the right, creating less fit single heterozygotes relative to the *abab* migrants coming from the left. This implies that on average the allele *a* is favored by selection and locally increases in frequency from the left, resulting in the *A* cline moving to the right. Symmetrically, allele *B* is favored by selection from the right, moving the *B* cline to the left. Without recombination, eventually the two clines meet and form double heterozygotes due to the presence of both *ab* and *AB* haplotypes. Even though the *Ab* haplotype exists before the alignment of clines and is the more fit single heterozygote, selection will disfavor both these single heterozygotes and double heterozygotes, yet is unable to remove the double heterozygotes as they continually form through mating of *ab* and *AB* haplotypes. In constrast, when we allow for recombination, the *ab* haplotype coming from the left will recombine locally and allow for the continual creation of single heterozygote individuals with genotype *abaB*. In this case, the selective advantage of allele *a* is slightly reduced on this genomic background that now exists, resulting in the clines moving slower towards each other as compared to the more rapid alignment of clines in the absence of recombination. This also leads to the observation that the clines in the presence of recombination are shallower than the steep clines that exist without recombination, creating a wider region of low fitness and overall greater load at the cline center when recombination occurs (Fig. S3).

Fig. 4 investigates the time it takes until two clines spatially align as a function of their initial distance, the selection intensity, the diffusion parameter, and the recombination rate. We find that clines will always couple, but over longer time spans the higher recombination is or the greater the initial distance between clines. The time it takes for clines to spatially align increases exponentially with increasing re-scaled initial distance (for a given *r* and a given *s*). Furthermore, for a given re-scaled initial distance, the time it takes for the clines to align is inversely proportional to the selection coefficient *s*, and the strength of attraction also rapidly declines if the re-scaled recombination rate *r*/*s* increases.

**Figure 4:**
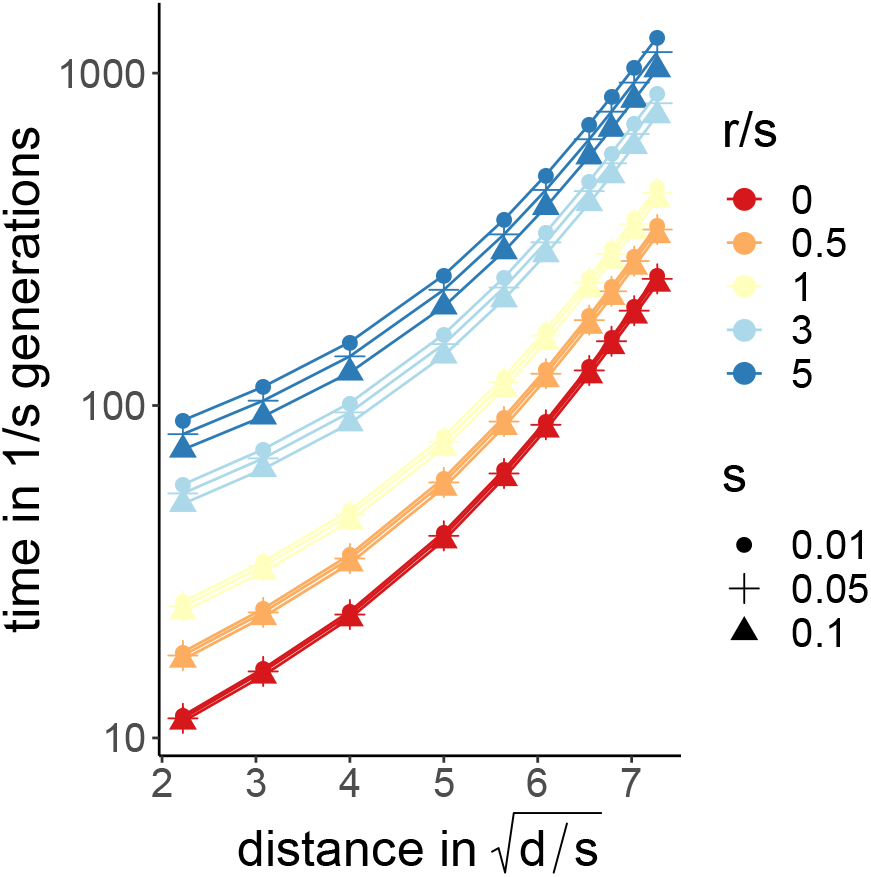
Time until clines coincide spatially, as a function of the re-scaled initial distance. The clines are initially vertical, i.e. the space is divided into three compartments where only one haplotype is present: *ab* to the left, *Ab* in the centre, and *AB* to the right. *s* is selection, *r* is recombination, and *d* is the diffusion parameter (see main text).

The patterns described above result because the clines described by our differential equations have an infinite spatial width, therefore even when they are far apart, there is always a partial overlap eventually leading to alignment of the clines. Our differential equations ignore genetic drift, creating important differences in behavior as compared to finite-sized populations. For finite populations, there should exist a critical distance between clines below which they attract each other and eventually align, and above which they remain independent due to non-overlap. To estimate this critical distance, we can approximate that the differential equation correctly describes clines in finite populations for frequencies in the range [*ϵ*, 1 *−ϵ*], and that outside of this range the allele is either fixed or lost. We take *ϵ* = 1/2*K*, expressing the fact that genetic drift dominates when a single copy of the allele is present in the deme. For a carrying capacity *K* = 50, we find that the effective spatial extent of a cline is 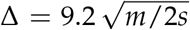 demes, where *m* is the migration rate of the discrete space model. If the clines are separated by a distance larger than Δ they do not overlap, and consequently they will not attract each other. If they are closer than Δ, they will potentially attract each other if selection is strong but recombination is weak, as observed in Fig. 4. For the parameters investigated in our forward simulations (Fig. 2, left panel, *m* = 0.25, *hs* = 0.1) we expect a critical distance of Δ = 10 demes between clines below which we would expect attraction and eventual cline alignment to occur. We observe, however, that many clines reach closer distances without aligning (e.g. see Fig. S2), indicating that genetic drift plays an important role even at close distances.

So far we have only considered clines that are neutral in the sense that both homyzgotes have the same fitness. Fig. 5 shows an example with one neutral cline (solid red lines) and a cline where the derived homozygote has a lower fitness than the wild-type (solid black lines). Thus, one cline moves towards the other one due to the selective advantage of the wild-type homozygotes. With strong recombination (*r* = 0.5, top panels) we find that the clines move past each other without much interaction. If the two clines are tightly linked (*r* = 0, bottom panels) we find that the two clines align and remain spatially aligned as they travel through space together.

**Figure 5:**
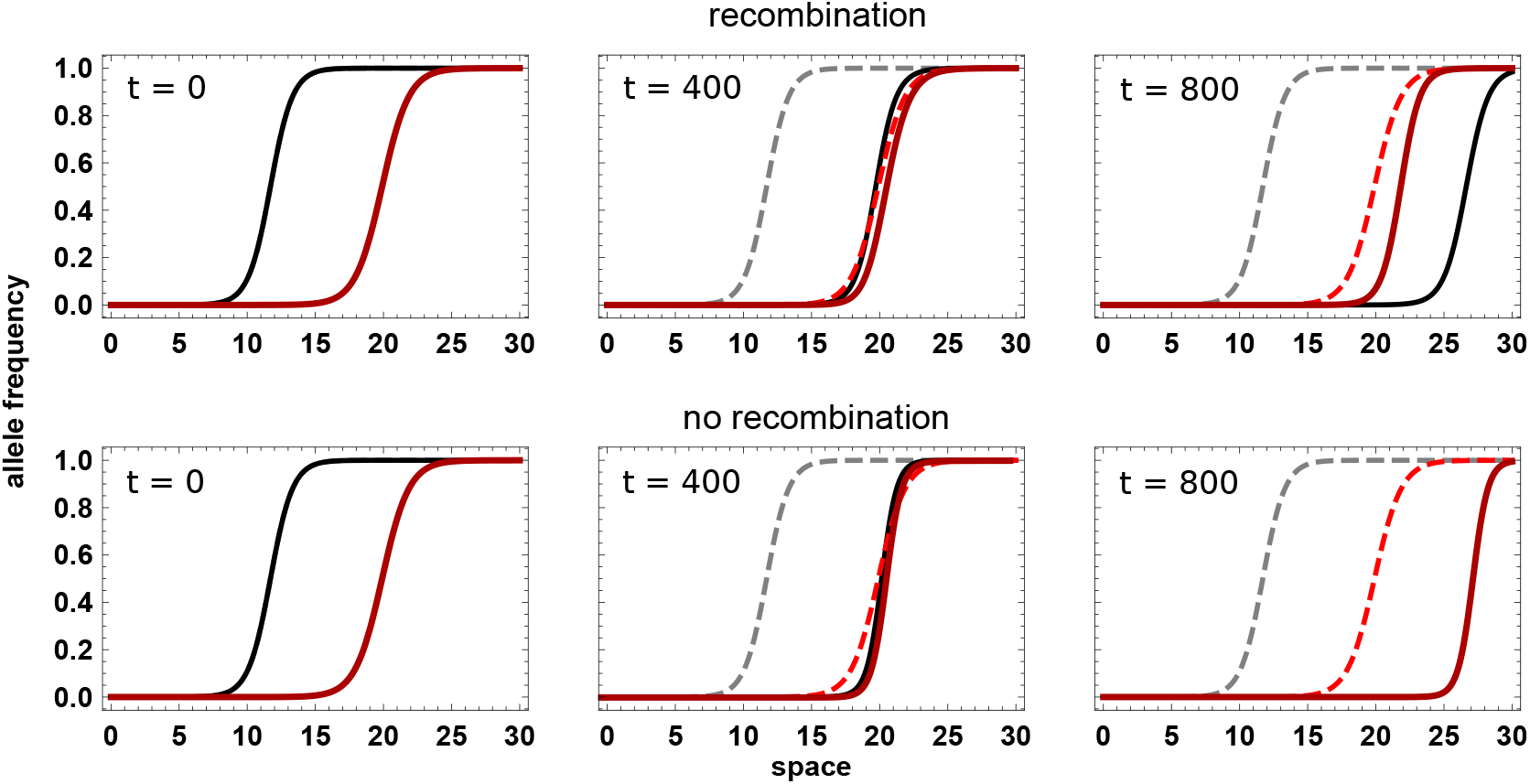
Temporal evolution of two clines at two different loci and spatial locations. At one locus (solid black lines) derived homozygotes have fitness 1 *− s*_*A*_ such that the cline moves through space from left to right. Parameter values: *s* = 0.1, *m* = 0.05, *s*_*A*_ = *−* 0.05 and (top row) *r* = 0.5 and (bottom row) *r* = 0.

## Discussion

In this study we have examined a little-studied phenomenon: the surfing of under-dominant alleles during range expansion and the subsequent movement over space of clines in allele frequencies established during surfing events. We derived the probability that underdominant mutations successfully establish at the expansion front in a serial founder event model (see Fig. 1). Unlike deleterious mutations that can lead to a transient increase in mutation load in recently colonized areas (Peischl et al., 2013), underdominant mutations will be maintained in the form of clines after a surfing event (see Fig. 2). Using individual-based simulations and mathematical modelling we have shown that once clines of disadvantageous heterozygotes form, clines can align in space due to differences in fitness on different genomic backgrounds and have the potential to combine as double (or more) heterozygotes. Our results show that the spatial alignment of clines is a stable state, corroborating recent theoretical results on the local stability of already aligned two-locus clines (Alfaro et al., 2021).

We have identified two main factors determining whether and how fast clines will spatially align: (i) clines need to be sufficiently close in space and (ii) the recombination distance between the two clines needs to be sufficiently low (see Figs. 3 & 4). An intuitive interpretation of these results is that clines will only attract each other if there is a spatial overlap of unfit single-locus heterozygotes such that unfit double heterozygotes are produced (see Fig. S3). When this overlap occurs, clines will move towards each other since the double homozygotes to either side of both clines are more fit than the single heterozygotes interior to the two overlapping clines. In the absence of recombination, once double heterozygotes are formed, they cannot be broken apart so selection cannot remove these genotypes at the cline center and a steep cline exists of low fitness as selection maintains double homozygotes on either side. Under high recombination, however, single heterozygotes are continually formed. Selection acts less strongly against single heterozygotes since they have higher fitness than double heterozygotes, explaining the slower movement and eventual alignment of clines in the presence of recombination (as well as the slightly wider region of space exhibiting lower fitness; see Fig. S3). The strength of selection has opposing effects on the spatial alignment of clines. Strong selection makes clines steeper (by decreasing the spatial scale 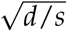), such that they become less likely to overlap. But strong selection also reduces the efficacy of recombination in creating fitter genotypes because multi-locus heterozygotes are efficently removed from the population which renders recombination ineffective. The resulting linkage disequilibrium in turn increases the strength of attraction of clines.

There are several important caveats and future directions to pursue related to our results. First, because our analytical model is based on diffusion in space, clines will always overlap and thus our model always predicts that clines will eventually merge (Fig. 4). However, in finite populations, clines will not extend indefinitely and genetic drift will play a larger role. We observe this when examining the movement of clines in our forward simulations (Fig. 2, right panel) and from the smaller observed critical distance (Fig. S2) below which two clines are predicted to merge, which is sensible as drift should reduce cline width as compared to deterministic expectations (Polechová and Barton, 2011). Second, we modeled simple single mutations that confer heterozygote disadvantage, yet there are a wide range of genomic or environmental scenarios that can cause heterozygote disadvantage as well as potential epistatic interactions. Genomic inversions may be the most prevalent example where heterozygotes are disfavored due to the suppression of recombination in the inverted region (Kirkpatrick, 2010). Polymorphism, and even ancient persistence, of inversions is widely observed in species across the globe (Hoffmann et al., 2004). Our model provides one potential explanation for the existence of such clines in inversion frequencies. We have also ignored the role of epistatic interactions between mutations at different loci. Our results suggest that any fitness landscape where multi-locus heterozygotes suffer a disproportionate fitness disadvantage a compared to single locus heterozygotes should hasten the spatial alignment of underdominant clines.

We have not investigated in depth the impact of unbalanced fitness between homozygotes in our model. Our clines, if alone, would remain stable and only fluctuate over space due to genetic drift. However, if either the derived or ancestral homozygotes are more fit than the other (with heterozygotes still being least fit), such clines would move over space after they have formed due to selection, until the less fit allele is purged from the population (Fig. 5). Furthermore, if the fitness advantage is confined to a certain region of the species range, the movement of the cline will stop at some point and form a stable tension zone (Barton and Hewitt, 1985; Bierne et al., 2011). Intriguingly, our model shows that if such moving clines merge with a neutral non-moving cline, these two clines will continue to travel together when recombination is sufficiently weak (Fig. 5). This suggests that underdominant alleles that confer a fitness advantage as homozygotes in a region of a species range, or beneficial alleles that are linked to underdominant alleles, may move through space and collect neutral clines that would otherwise not move, reinforcing genetic differentiation across aligned clines.

The role of range expansions on the establishment and subsequent alignment of clines in space has important ramifications, since over evolutionary timescales such clines will be expected to couple with more clines and reinforce more strongly any reproductive barrier one clines already imposes (Butlin and Smadja 2018, see also Alfaro et al. 2021). It will be fruitful to pursue empirical investigations into the prevalence of underdominance in regions of partial reproductive isolation across species ranges. Since heterozygote disadvantage does not necessitate genetic underpinnings, even the “surfing” of cultural behavior, such as learned birdsong could equally contribute regions of heterozygote disadvantage arising during range expansions. Previous simulation work on ring species has shown the instability of species barriers which form but will dissipate after 10s of thousands of generations (Martins et al., 2013). Our results, however, suggest that such formation of barriers between species may be even more stable, as aligned clines of heterozygote disadvantage should persist through time. These phenomena could potentially play a large role in the process of speciation as pre- and post-zygotic barriers are reinforced at a localized area of a species range.

## Acknowledgments

We would like to thank Nick Barton, Laurent Excoffier, and Jitka Polechova for useful feedback while working on this project. KJG was funded by Swiss National Science Foundation Ambizione grant PZ00P3 _185952.

## Appendix

### A Derivation of establishment probability

We use a diffusion approach to calculate the fixation probability of underdominant mutations in a single panmictic population of size *N*. We consider a single locus with two alleles *a* and *A*. Fitness is given by *w*_*aa*_ = 1, *w*_*aA*_ = 1 *− s* and *w*_*AA*_ = 1. Let *p* denote the frequency of allele *A*. The variance in allele frequency change due to binomial sampling is

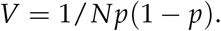

The expected change in frequency of allele *A* due to selection is given by

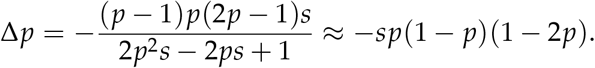

Using standard diffusion approximation results, the probability of fixation (conditioned on initial frequency *p*_0_) is given by:

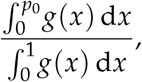

where

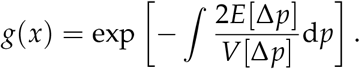

The solution for the fixation probability can be expressed using the error function:

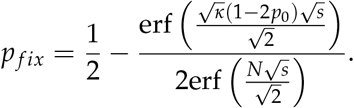

For weak selection this can be further approximated by

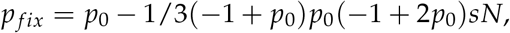

We can thus define an effective selection coefficient

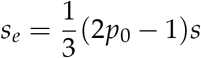

which we use as the selection coefficient in Kimura’s original equation for the fixation probability such that we can write 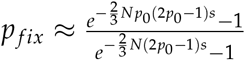.

In the asymmetric case analogoues calculations yield

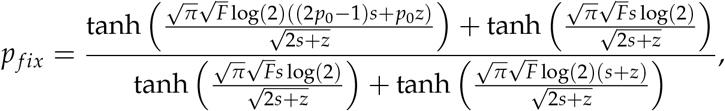

where we have used 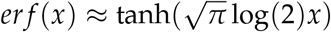.

We note that a relatively simple weak selection approximation can also be derived in the asymmetric case but we found that this approximation is in general not very accurate.

For the serial founder model, we use a heuristic approach motivated by the derivations in (Peischl et al., 2013, 2015) and simply replace *s* with *sT* and *N* with *F*. Fig. 1 in the main text shows a comparison of individual-based simulations and the heuristic approximation and reveals that the approximation indeed works reasonably well.

### B Additional Figures

**Figure S1:**
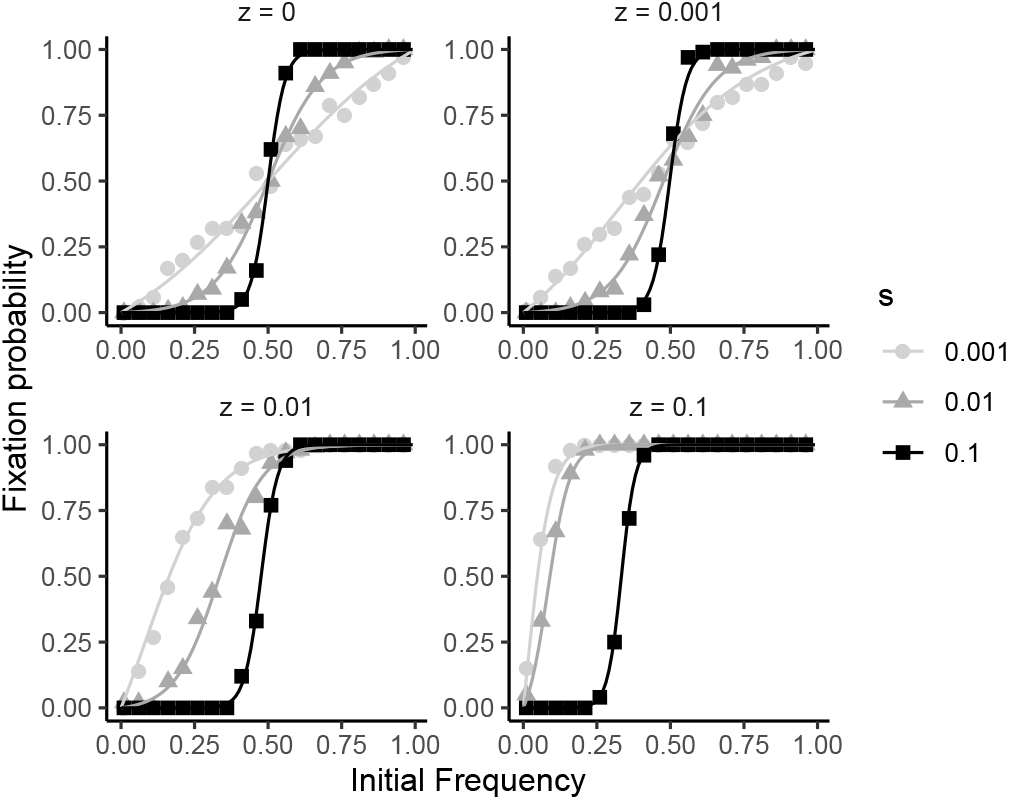
Fixation probability of an underdominant, beneficial mutation in a population of constant size (*N* = 500 diploid individuals). The figures show the fixation probability of a mutation *A* as a function of its initial frequency. The fitness scheme is *w*_*aa*_ = 1, *w*_*aA*_ = 1 *− s*_1_, *w*_*AA*_ = 1 + *s*_2_

**Figure S2:**
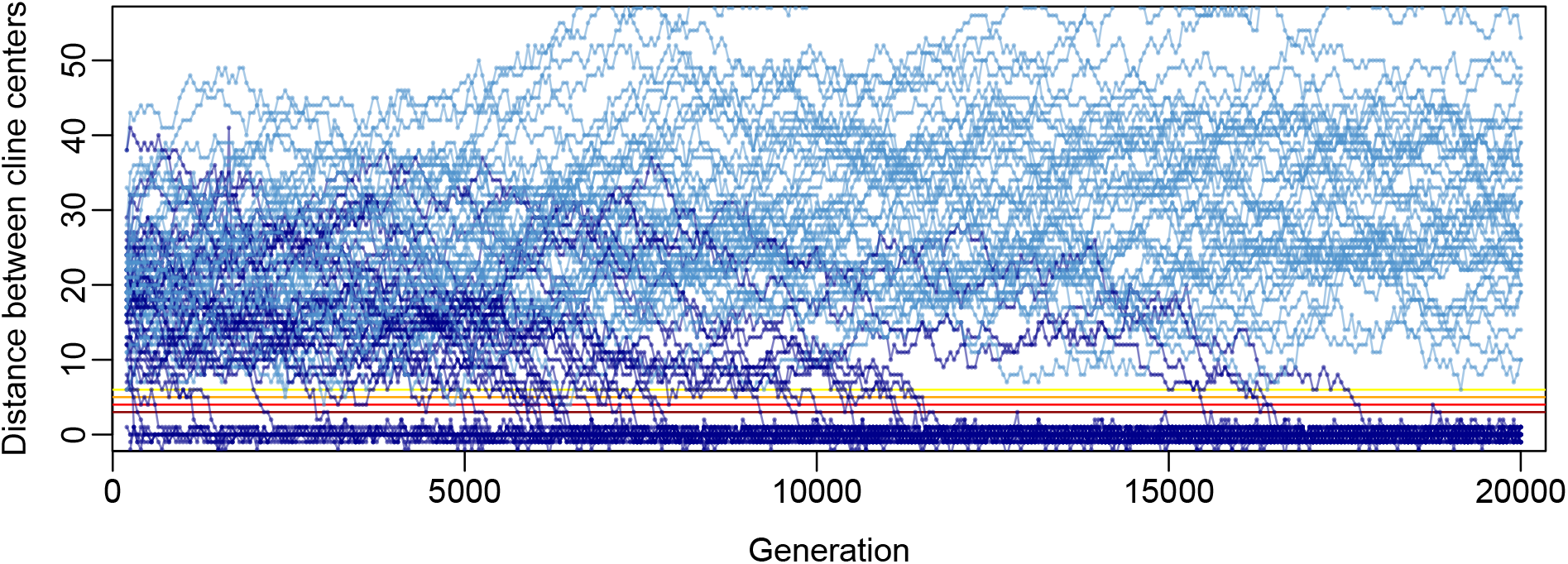
The distance between adjacent cline centers plotted through time. Because cline centers are measured as the deme with allele frequency closest to 0.5, but clines are steep, there is a +/ *−* 1 deme margin in accuracy for the distance between cline centers, hence some *−* 1 distance values. Dark blue lines are those cline centers that eventually couple, while light blue lines are those cline centers that never couple across the course of the simulations. These simulations are the same as in Fig. 2, with 5 manually introduced underdominant alleles, *m* = 0.25, *K* = 50, and heterozygote fitness is 0.9 for single-mutant heterozygotes. 20 replicate simulations with 5 loci each are shown.

**Figure S3:**
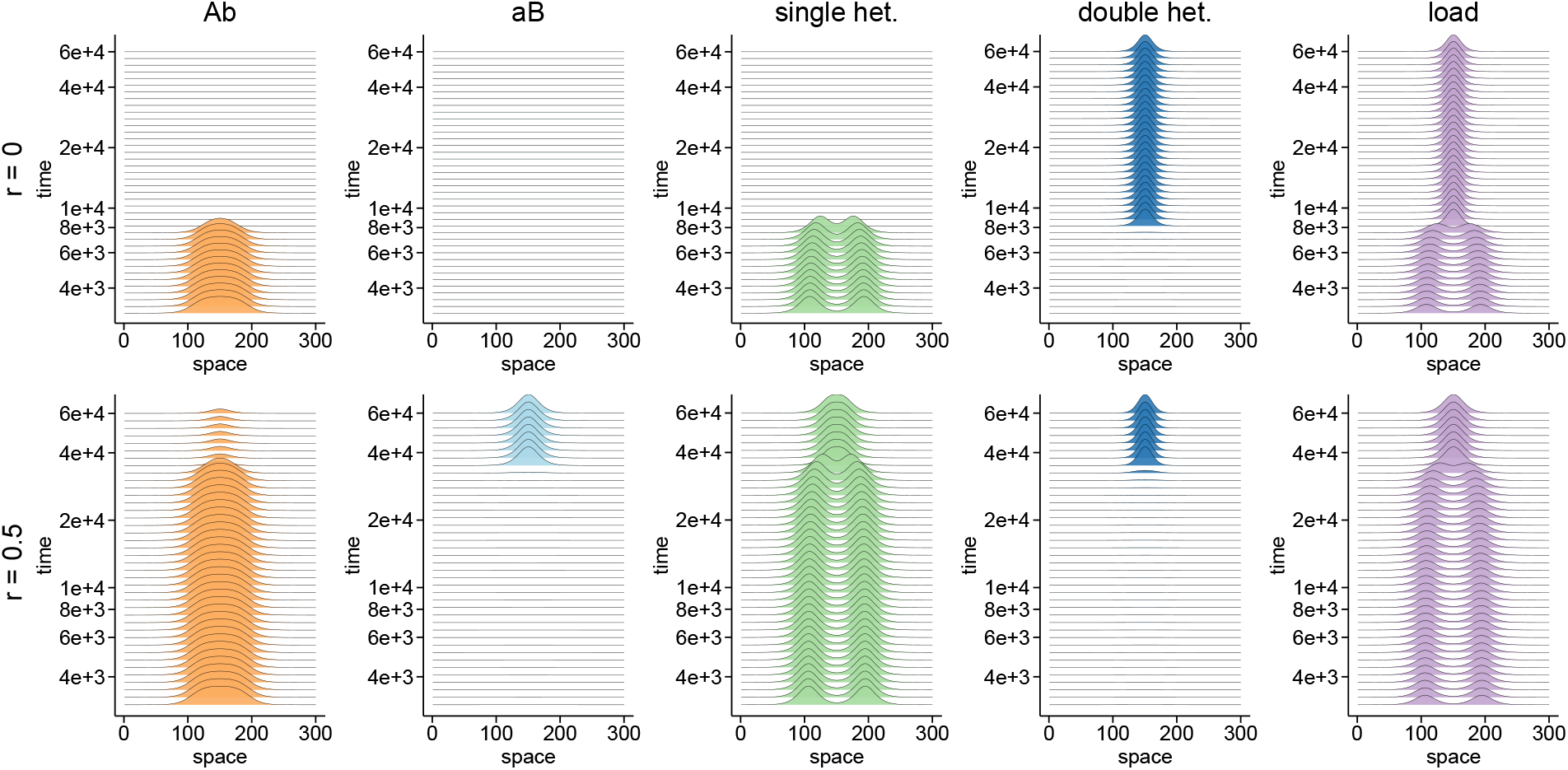
The evolution of haplotype frequencies and genetic load during the course of two clines attracting and then aligning, from differential equation 2. The top row shows results for no recombination and the bottom row shows results for free recombination. The landscape consists of 300 demes (x-axis), and parameter values are *s* = 0.1 and *d* = 0.05. At *t* = 0 the clines are vertical and are located at demes 105 and 195. One deme corresponds to 0.095 *x*_0_ spatial units. Over time (generations on the y-axis), the two clines are attracted to each other and eventually align (time *t* = 7896 for *r* = 0, and *t* = 33781 for *r* = 0.5) at the central deme, 150, of the landscape. In the case of no recombination, haplotype *aB* never exists since the *bB* cline exists to the right of the *aA* cline, so the derived *B* allele can never recombine onto the ancestral *a* background. In this case, once clines merge, all individuals are either double homozygotes or double heterozygotes across the entire landscape, creating a narrow range of high load at the cline centers. Conversely, with recombination, both single heterozygote haplotypes exist, and after the clines align, both double and single heterozygotes persist over a slightly wider deme width around the cline center, since the cline with recombination is shallower.

